# Molecular mechanism of sulfur chemolithotrophy in the betaproteobacterium *Pusillimonas ginsengisoli*

**DOI:** 10.1101/708438

**Authors:** Subhrangshu Mandal, Moidu Jameela Rameez, Prosenjit Pyne, Sabyasachi Bhattacharya, Jagannath Sarkar, Wriddhiman Ghosh

**Author notes:** National Institute of Cholera and Enteric Diseases(NICED), P- C.I.T. Scheme XM, Beleghata, 33, CIT Rd, Beleghata, Kolkata - 700054, India. **Correspondence emails** /.

## Abstract

Molecular mechanism of chemolithotrophic sulfur oxidation in *Betaproteobacteria* is less explored than that in *Alphaproteobacteria*. Here we carried out whole genome sequencing and analysis of a new betaproteobacterial isolate *Pusillimonas ginsengisoli* SBSA which oxidizes thiosulfate via formation tetrathionate as an intermediate. The 4.7-Mb SBSA genome was found to encompass a complete *soxCDYZAXOB* operon, plus one thiosulfate dehydrogenase *(tsdA)* and sulfite:acceptor oxidoreductase (*sorAB*) genes. Recombination-based knock-out of *tsdA* revealed that the entire thiosulfate oxidized by SBSA is first converted to tetrathionate, and no thiosulfate is directly converted to sulfate as typical of the Alphaproteobacterial Sox pathway whereas its tetrathionate-oxidizing ability was as good as that of the wild-type. The ∆*soxYZ* knock-out mutant exhibited wild-type-like phenotype for thiosulfate/tetrathionate oxidation, whereas ∆*soxB* oxidized thiosulfate only up to tetrathionate and had complete impairment of tetrathionate oxidation. However, substrate-dependent O_2_-consumption rate of whole cells, and sulfur-oxidizing enzyme activities of cell-free extracts, measured in the presence/absence of thiol-inhibitors/glutathione, indicated that glutathione plays a key role in SBSA tetrathionate oxidation. All the present findings collectively indicated that glutathione:tetrathionate coupling in *Pusillimonas ginsengisoli* may involve some unknown proteins other than thiol dehydrotransferase(ThdT), while subsequent oxidation of the potential glutathione:sulfodisulfane and sulfite molecules produced may proceed via *soxBCD* action.

## 1. Introduction

Awing to a wide range of oxidation states (2-to 6+) the element sulfur enters into a various biogeochemical processes. Lithotrophic oxidation of reduced sulfur compounds is considered as an ancient metabolism that, in course of evolution, has mechanistically diversified in different groups of *Bacteria* and *Archaea* [1, 2] Of the major biochemical pathways for sulfur-chemolithotrophy known thus far [3–7], the one governed by the Sox multi-enzyme system (SoxXAYZBCD) is the best-studied. This pathway, typically described from members of the class *Alphaproteobacteria*, is known to directly oxidize sulfide, elemental sulfur, thiosulfate or sulfite (but none of the polythionates) to sulfate, without the formation of any free intermediate [3, 8–14]. Albeit *soxXAYZBCD* genes are widespread in phylogenetically-diverse bacteria, no chemolithotroph outside the class *Alphaproteobacteria* has so far been reported as using the typical Sox mechanism for sulfur oxidation. Only a few strains of *Guyparkeria* under the class *Gammaproteobacteria* oxidize thiosulfate directly to sulfate via what appears to be a Sox-like mechanism [15]. In contrast, almost all other beta- and gammaproteobacterial chemolithotrophs oxidize thiosulfate via formation of tetrathionate intermediate (S_4_I) and subsequent oxidation of the latter to sulfate [5, 16–21].

So far as the S_4_I pathway is concerned, mechanism of the sulfur chemolithtrophy described mostly in obligate extremophiles where thiosulfate dehydrogenation (tetrathionate formation) and tetrathionate oxidation are rendered by Archaeal homologs TQO and TetH respectively. This pathway with restricted phylogenetic spread in bacteria, does not however explains the mechanism involved in S_4_I pathway encountered in the facultative counterparts [1, 5, 22–23]. In contrast to the above scenario, molecular mechanism of S_4_I pathway in facultative sulfur chemolithotrophs are less explained. Recently, a novel S_4_I pathway of thiosulfate oxidation has been reported from the betaproteobacterium *Advenella kashmiren*sis [24] where thiosulfate-to-tetrathionate conversion is carried out by the action of a thiosulfate dehydrogenase related to the TsdA protein epitomized in the anoxygenic photolithotroph *Allochromatium vinosum* [25, 26] as well as a novel protein thiol dehydrotransferase (ThdT). The ThdT, in *A*. *kashmirensis*, additionally catalyzes glutathione-mediated tetrathionate oxidation [24].

Here we reveal the molecular mechanism of sulfur-chemolithotrophy in the *Pusillimonas ginsengisoli* strain SBSA (MTCC12558) isolated from a sediment-sample retrieved from 275 centimeters below the sea-floor (cmbsf), at a water-depth of 530 meters below the sea-level (mbsl), of the Eastern Arabian Sea off the west coast of India. The whole-genome of *P*. *ginsengisoli* SBSA was sequenced and analyzed to identify gene loci encoding potential sulfur oxidation proteins, while role of putative sulfur oxidation genes were investigated by carrying out knock-out mutations and testing the chemolithotrophic abilities of the resulting mutants. Physiological and biochemical studies were carried out to corroborate the conclusions derived from the genomic/genetic data. Results showed that some of the enzymes used by *Pusillimonas ginsengisoli* SBSA to oxidize thiosulfate and tetrathionate are similar to those of *A*. *kashmirensis*, whereas some of them are potentially distinctive.

## 2. Materials and methods

### 2.1 Isolation and characterization of *Pusillimonas ginsengisoli* strain SBSA

During the research cruise on-board RV Sindhu Sankalp (SSK42), one ∼3-m-long gravity core numbered as SSK42/6 was collected and sampled as described previously, from 530 mbsl water-depth of the Eastern Arabian Sea, off the west coast of India (16°50.03’ N, 71°59.50’ E) [27]. Sediment-sample from 275 cmbsf of this core was used to isolate aerobic sulfur chemolithotrophs via 14-day enrichment (5% w/v) in modified basal and mineral salts (MS) solution [28] supplemented with sodium thiosulfate (20 mM Na_2_S_2_O_3_.5H_2_O) and 500 mg yeast extract L^−1^ (MSTY). The sediment-MSTY mixture was incubated at 15°C on a rotary shaker until the phenol red indicator of the medium turned yellow. Following this, strain SBSA was isolated from the mixture, alongside several other neutrophilic, mesophilic and facultatively sulfur-chemolithotrophic bacteria, via serial dilution plating, and subsequent dilution streaking, on MSTY-agar plates. Besides the mixotrophic MSTY medium, the new isolate was found to grow and produce sulfate in mixotrophic MSTrY medium (MS-tetrathionate-yeast extract; 10 mM K_2_S_4_O_6_ and 500 mg yeast extract L^−1^ medium); and was maintained in chemoorganoheterotrophic i.e. Luria–Bertani (LB) medium. SBSA however did not grow in chemolithoautotrophically in MST or MSTr medium that lacked yeast extract. The thiosulfate and tetrathionate oxidizing chemoorganoheteroph SBSA was taxonomically classified down to the lowest identifiable taxonomic category, as described previously [19], on the basis of their 16S rRNA gene sequence similarities with validly-published species (http://www.bacterio.net/) [29, 30].

### 2.2 Physiology experiments

Relevant bacterial strains and plasmids used in this study are listed in Table 1. All strains, wild-type or engineered, were grown and maintained in Luria-Bertani (LB) broth or agar-plates, which were supplemented with suitable antibiotic(s) whenever needed. Chemolithoautotrophic properties of the *P. ginsengisoli* wild-type, or its different mutants, were tested in MSTY/MSTrY medium. For all experiments in MS-based media, seed cultures of *P. ginsengisoli* wild-type/mutants were prepared in LB broth; 1% (v/v) inocula were then transferred from mid-log phase seed-cultures to experimental cultures in MS-based media. *E. coli* were incubated aerobically at 37°C, whereas *P. ginsengisoli* and its mutants were incubated aerobically at 28°C. Broth cultures were kept in incubator-shakers at 150 × g. Chemolithotrophic oxidation potential of reduced sulfur compounds was checked in MSTY or MSTrY media correspondingly. For this purpose, 5 ml cultures were taken at appropriate time points, centrifuged at 9500 × g for 5 min, and concentrations of dissolved thiosulfate, tetrathionate and sulfate in the supernatant finally measured by iodometric titration, cyanolytic method and gravimetric precipitation method, respectively [19, 20, 31, 32]; sulfite was measured spectrophotometrically with pararosaniline as the indicator [33]. To corroborate the results obtained by the above methods, sulfate, sulfite and thiosulfate were further quantified by anion chromatography via chemical suppression, using an Eco IC (Metrohm AG, Switzerland) equipped with a conductivity detector (Metrohm, IC detector 1.850.9010) and autosampler (863 Compact Autosampler, Switzerland). Separation of ions was carried using a Metrosep A Supp5 - 250/4.0 (6.1006.530) anion exchange column (Metrohm AG); a mixed solution of 1.0 mM sodium hydrogen carbonate and 3.2 mM sodium carbonate was used as the eluent; 100 mM sulfuric acid was used as the regenerant; flow rate was 0.7 mL min ^−1^, and injection volume 100 µL. Samples were diluted 1000-fold with de-ionized water (Siemens, <0.06 μS), and filtered by passing through 0.22 µm hydrophilic polyvinylidene fluoride membranes (Merck Life Science Private Limited, India), prior to analysis. Sigma-Aldrich (USA) standard chemicals were used to prepare the calibration curve for quantification. Overall sample reproducibility was ±0.2 ppm. To measure substrate-dependent oxygen consumption, cells grown in MSY, MSTY or MSTrY were harvested from mid-log phase cultures grown in MSY, while harvesting of cells from MSTY- or MSTrY-grown cultures was done when 80% of the supplied thiosulfate had been converted to tetrathionate or 80% of the supplied tetrathionate was converted to sulfate respectively. Harvested cells were washed twice with 50 mM potassium phosphate buffer (pH 7.5) and finally resuspended in the same buffer. Cells were maintained for no longer for 2 h at 30^°^C before being used for oxygen consumption assays. Oxidation of reduced sulfur compounds was thenmeasured polarographically in a Biological Oxygen Monitor equipped with a Clarke-type oxygen electrode (Yellow Springs Instruments, Yellow Springs, OH, USA), at 30^°^C, in potassium phosphate buffer (pH 7.5) with cells equivalent to 300 mg of proteins in 3 ml reaction volumes. Reaction mixtures contain following substrate concentrations: sodium thiosulfate 10 mM; potassium tetrathionate 5 mM and sodium sulfite 2 mM in 5 mM EDTA. Calculations were based on the assumption (according to the instrument manual) that an 18% change in oxygen concentration equivalent to 2.71 ml of oxygen consumed. In all the above cases, oxygen consumption rates (expressed as nmol of oxygen consumed mg cellular protein^−1^ min^−1^) were corrected for chemical- or auto-oxidation of substrates and endogenous respiration rates. For verifying the potential role of thiol groups in substrate dependent oxygen consumption rates of SBSA, harvested cells were treated with the thiol-binding reagents NEM or iodoacetamide (1 mM of the tested reagent for 10 min), following which their oxygen consumption rates were determined and compared with the rates determined for untreated cells. For NEM- or iodoacetamide-treated cells, oxygen consumption rates were additionally measured in potassium phosphate buffer having pH 8.5 [since these reagents react optimally with the thiolate anion resulting from deprotonation of thiol group (typical pKa 8.3) [34], over and above the tests in the potassium phosphate buffer having pH 7.5. To eliminate the excess NEM or iodoacetamide before polarographic assays, inhibitor-treated cells were then washed with 50 mM potassium phosphate buffer having pH 7.5 or 8.5, according as the pH of the assay buffer was 7.5 or 8.5. Notably, substrate-dependent oxygen consumption rates of inhibitor treated cells did not change during changing the pH of the assay buffer to 8.5 from 7.5. Therefore, only the data obtained by using the assay buffer of pH 7.5 are shown in Fig. 4A–C. Potential effect of GSH on the sulfur substrate dependent oxygen consumption by SBSA was further verified individually by treating the harvested cells with 10 mM GSH for 10 min prior to the oxygen consumption assays. Treated cells were washed with 50 mM potassium phosphate buffer (pH 7.5) to get rid of extra GSH. Oxygen consumption values obtained in this way for GSH-treated sets were finally compared with the values already obtained for the untreated ones.

**Table 1.**
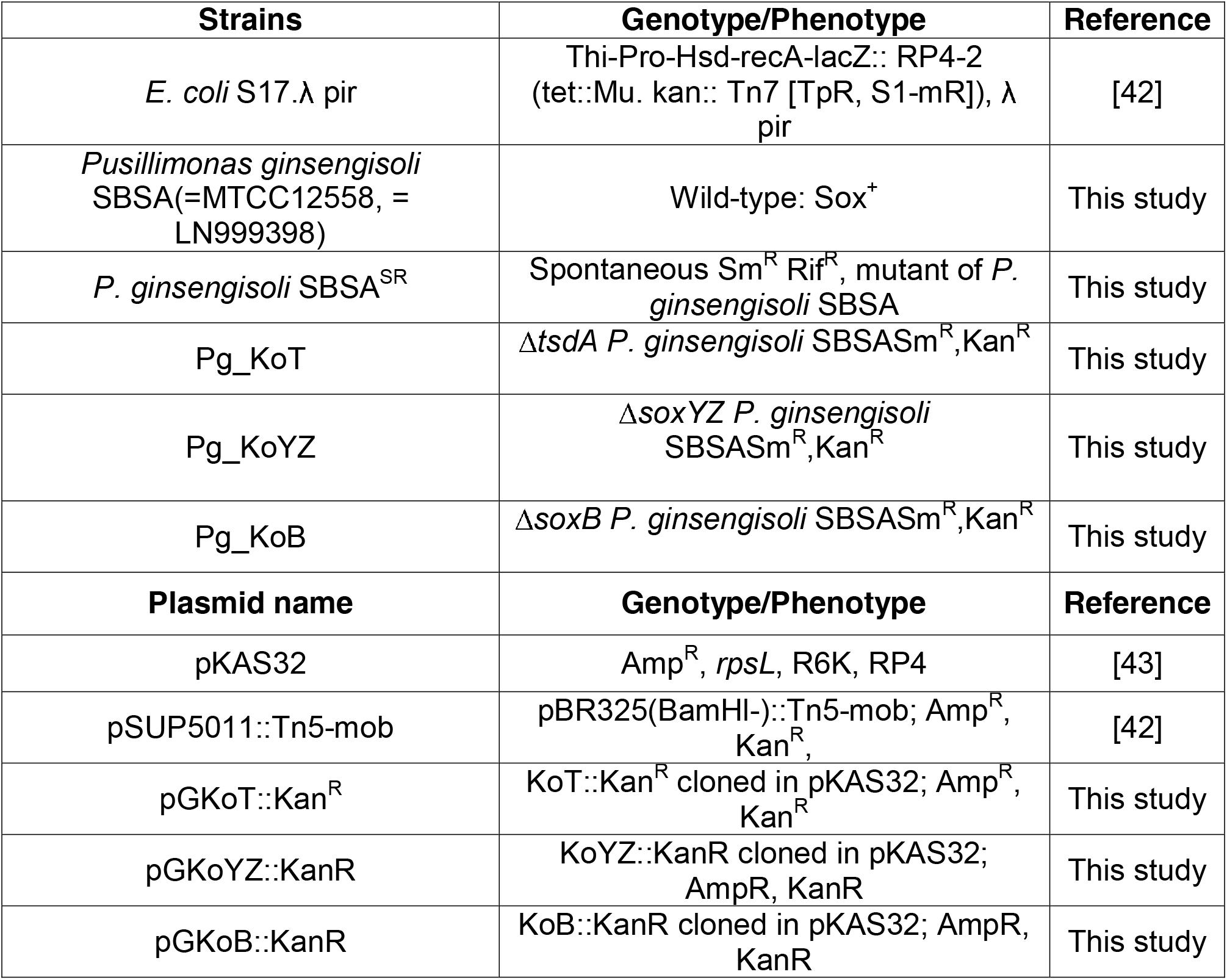
Bacterial strains and plasmid used in this study.

### 2.3 Biochemical experiments

Activities of the thiosulfate-, tetrathionate- or sulfite oxidizing enzyme systems were assayed in the CFEs of MSY-, MSTy- or MSTry-grown cultures. Cells were were harvested at the same growth stages as stated for the substrate dependent oxygen consumption assays. Harvested cells were washed twice and resuspended in 50 mM potassium phosphate buffer (pH 7.5), following which they were lysed by sonication. After high-speed centrifugation at 16,000 × g for 20 min, supernatants were taken out in fresh tubes as crude CFEs and their protein concentrations determined by Bradford method. Thiosulfate dehydrogenase activity in the CFEs was determined by a method improvised from that described earlier [35]. A reaction mixture (2.0 ml) containing 50 mM potassium phosphate buffer (pH 7.5), 2 mM potassium ferricyanide, 7.5 mM of sodium thiosulfate and CFE (≈200 µg of protein) was prepared to start the reaction. Enzyme activity was determined spectrophotometrically by measuring the rate of ferricyanide reduction at 420 nm. Activity of the tetrathionate-oxidizing enzyme system in the crude CFEs was then estimated according a method similar to the one used for thiosulfate dehydrogenase, except for the fact that here 7.5 mM potassium tetrathionate was added instead of sodium thiosulfate. Sulfite dehydrogenase / oxidase activity in the CFEs was assayed using a method adapted from one described earlier [12]. Here, the reaction mixture (2.0 ml) contained 50 mM Tris-HCl (pH 7.9), 2 mM potassium ferricyanide, 3 mM sodium sulfite in 5 mM EDTA and CFE (≈200 µmg of protein). To verify the potential effect of glutathione on the activities of these enzyme systems, the above reactions were again carried out individually by adding 10 mM GSH to the respective reaction mixtures. Total enzyme activity and specific activity in all the above cases were finally expressed as µmol of ferricyanide reduced min^−1^ and µmol of ferricyanide reduced mg protein^−1^ min^−1^ correspondingly.

### 2.4 Whole genome sequencing and annotation

Genomic DNA of *P. ginsengisoli* SBSA isolated from stationary phase culture was sequenced using Ion Torrent Personal Genome Machine (Ion PGM) 318 chip with 400bp read chemistry. The 1500000 high-quality reads obtained from the Ion PGM run (mean read length 178.81 ± 41.53 bp) were assembled using SPAdes 3.13.0 [36] in single cell mode. This assembly yielded 63 contigs which constituted a consensus of 4761130 bp. Of these, 63 contigs that were > 200 bp long were annotated using the automated Prokaryotic Genome Annotation Pipeline (PGAP) of the National Center for Biotechnology Information (NCBI; Bethesda, MD, USA) and deposited at DDBJ/ENA/GenBank under the accession RAPG00000000, the version described in this paper is RAPG00000000.1. All information regarding this Whole Genome Shotgun project is available in the GenBank under the BioProject PRJNA492192, while the raw sequence reads obtained from the Ion PGM run are available in the NCBI Short Read Archive (SRA) under the run accession number SRS3804619.

Completeness of the *P. ginsengisoli* SBSA genome was determined using CheckM 1.0.12 [37]. For this, a database of marker genes specific for the family *Alcaligenaceae* was constructed from the taxonomic marker genes provided with CheckM. Homologs of these marker genes were searched in the assembled draft genome of SBSA using HMMER algorithm. Completeness of the SBSA genome was calculated on the basis of the number of *Alcaligenaceae*-specific markers, detected in the genome of SBSA.

Homologs of genes potentially involved in bacterial sulfur oxidation were then searched either manually in the PGAP-annotated SBSA genome or by mapping/aligning query gene sequences from other betaproteobacterial species onto the *P. ginsengisoli* SBSA genome sequence using the PROmer utility of the MUMmer 3.23 package [38]. Annotations of the ORFs identified by the latter method were corroborated by NCBI BlastP analysis of the putative amino acid sequences, followed by their conserved domain search with reference to the Conserved Domain Database (CDD).

### 2.5 Knock out mutagenesis

Homologous recombination-based techniques were used to generate insertional mutations in the *soxB* (D7I39_07195), *soxYZ* (D7I39_07170 and D7I39_07175) and *tsdA* (D7I39_17460) genes of *P. ginsengisoli* SBSA. Since the *tsdA*, *soxYZ* and *soxB* sequences does not contain any unique restriction site suitable for the insertion of an antibiotic resistant cassette, a new *Kpn*I site had to be incorporated into these genes via fusion PCR method [24] and a kan^R^ cassette, amplified from the plasmid pSUP5011::Tn5, was then inserted into this *Kpn*I site. All the primers which were used to generate this construct have been listed in Supplementary Table S1-S3. The final constructs PGKoT::Kan^R^, PGKoYZ::Kan^R^ and PGKoB::Kan^R^, cloned in pKAS32 (see supplementary information for detailed procedure of knock out mutagenesis), were transformed into *Escherichia coli* S17.1 λ*pir* and used as donor strains in conjugative plate mating for transferring corresponding gene replacement cassettes to *P. ginsengisoli* SBSA. A spontaneous mutant strain *P. ginsengisoli* SBSASR, which was resistant to rifampicin (50 µg mL^−1^) and streptomycin (100 µg mL^−1^) was used as the recipient in the conjugation experiments. Cells from mid-log phase cultures of both the donor and the recipient were harvested and washed twice and re-suspended in 10 mM MgSO_4_. Donors and recipients were then mixed at a 1:6 cell mass ratio and immobilized onto a membrane filter with pore size of 0.2 μm. The membrane filter was placed on an LA plate containing 1% agar, and incubated at 30^°^C for 18 h. Cell mass growing on the membrane was harvested and dissolved in 2 mL 0.9% NaCl solution; this was subsequently spread at 10^−1^ dilution on LA plates containing both rifampicin (50 µg mL^−1^) and kanamycin (40 µg mL^−1^). Recipients which had successfully exchanged their relevant genomic loci with gene replacement cassettes loaded on the constructs were screened for double cross-over mutants by positively selecting for the cartridge-encoded kanamycin resistance trait and loss of streptomycin sensitivity conferred by the *rpsL* gene of pKAS32. Whilst the resultants ∆*tsdA*, ∆*soxB* and *soxYZ*::Kan^R^ mutants were designated as Pg_KoT, Pg_KoB and Pg_KoYZ respectively, orientations of the antibiotic resistance cassette with respect to the transcriptional directions of the ORFs mutated were checked by PCR-amplification and sequencing of DNA fragments containing specific genome-cartidge intersections.

## 3. Results

### 3.1 Thiosulfate and tetrathionate oxidation potential of *Pusillimonas ginsengisoli* SBSA

From 275 cmbsf of SSK42/6, four such aerobic bacterial strains were isolated in MSTY media that could utilize thiosulfate/ tetrathionate as a chemolithotrophic substrate when grown in corresponding chemolithotrophic medium (Fig. 1B and C). 16S rRNA gene sequence-based taxonomic identification of the isolates clustered them under single species-level entities belonging to *Pusillimonas*. These isolates not only formed tetrathionate from thiosulfate but also oxidized tetrathionate to sulfate. Among the four strains, one strain was selected (on which the present study was based; SBSA =MTCC12558) for detailed analysis of the molecular mechanism of sulfur chemolithotrophy.

**Figure 1.**
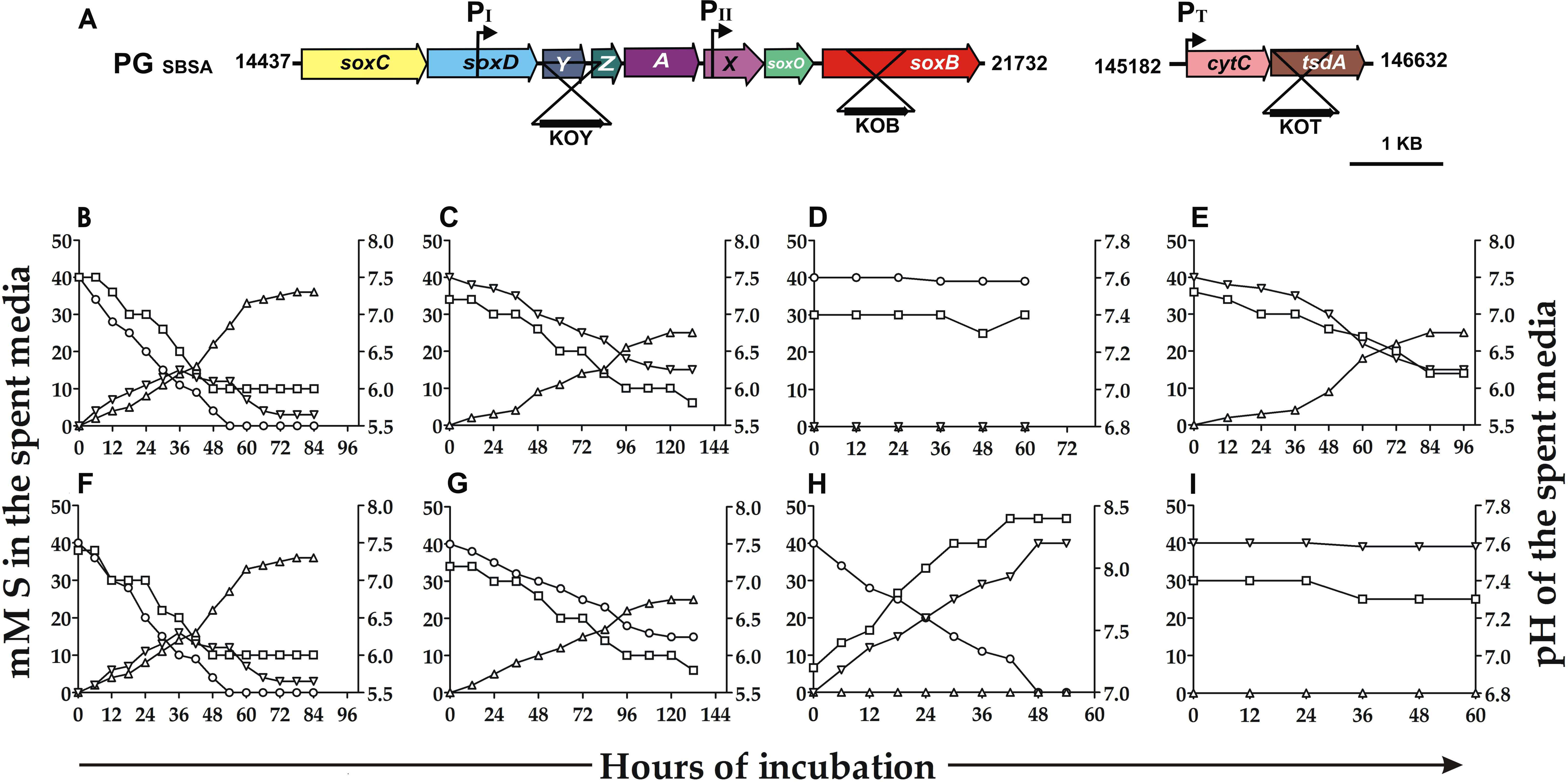
Homologous-recombination-based insertional mutation of the tsdA, *soxYZ* and *soxB* genes of *P. ginsengisoli*, and test of the sulfur-oxidation phenotypes of the resultant mutant strains. A. Physical map of the *sox* operon showing location of the potential promoters, plus the positions within the ORFs where antibiotic-resistance cassettes were inserted for mutant generation. Nucleotide numbers indicate the span of this operon within the *P. ginsengisoli* SBSA genome. Arrows indicate the orientation of the antibiotic-resistance cassettes. B and C. Kinetics of thiosulfate and tetrathionate oxidation by the wild-type strain SBSA respectively. D and E. Kinetics of thiosulfate and tetrathionate oxidation by the Δ*tsdA* mutant Pg_KoT respectively. F and G. Kinetics of thiosulfate and tetrathionate oxidation by the ∆*soxYZ* mutant Pg_KoYZ respectively. H and I. Kinetics of thiosulfate and tetrathionate oxidation by the ∆*soxB* mutant Pg_KoB respectively. -○-, - ▽- and -∆- and denote the concentration of sulfur (in mM S) in the spent media, at any time-point of incubation in the form of thiosulfate, tetrathionate and sulfate respectively. - □-denotes the pH of the spent medium at any given time point of incubation.

In MSTY medium (containing 40mM S thiosulfate as a chemolithotrophic substrate), simultaneous formation of tetrathionate and sulfate was observed from the first 6 h. At the 36^th^ h of incubation, 11 mM S of the supplied thiosulfate, 15 mM S tetrathionate and 14 mM S sulfate were detected in the spent medium. The supplied thiosulfate was completely extinct from the medium after an incubation period of 54 h while after a total incubation period of 72 h, 36 mM S sulfate and 3 mM S tetrathionate were remained in the spent medium (Fig1B). SBSA when grown in MSTrY medium (containing 40mM S tetrathionate as a chemolithotrophic substrate) able to oxidize 35 mM S tetrathionate into corresponding amount of sulfate over 120 h of incubation (Fig. 1C).

### 3.2 Genomic overview and sulfur oxidation related genes in *P.ginsengisoli* SBSA

Complete genome of *P.ginsengisoli* comprising a sequence consensus of 4,763,470 bp was obtained with an average read coverage of 92x. The average GC content of the genome was found to be 55.5% (Fig. 2.) while the highest contig size was 860965 bp. A completeness level of 99.16% was estimated for the assembled draft genome of SBSA (using the software CheckM) based on the presence of 483 out of a total 486 marker genes obtained from 27 representative genomes under the family *Alcaligeneaceae*; curated in the CheckM database.

**Fig. 2.**
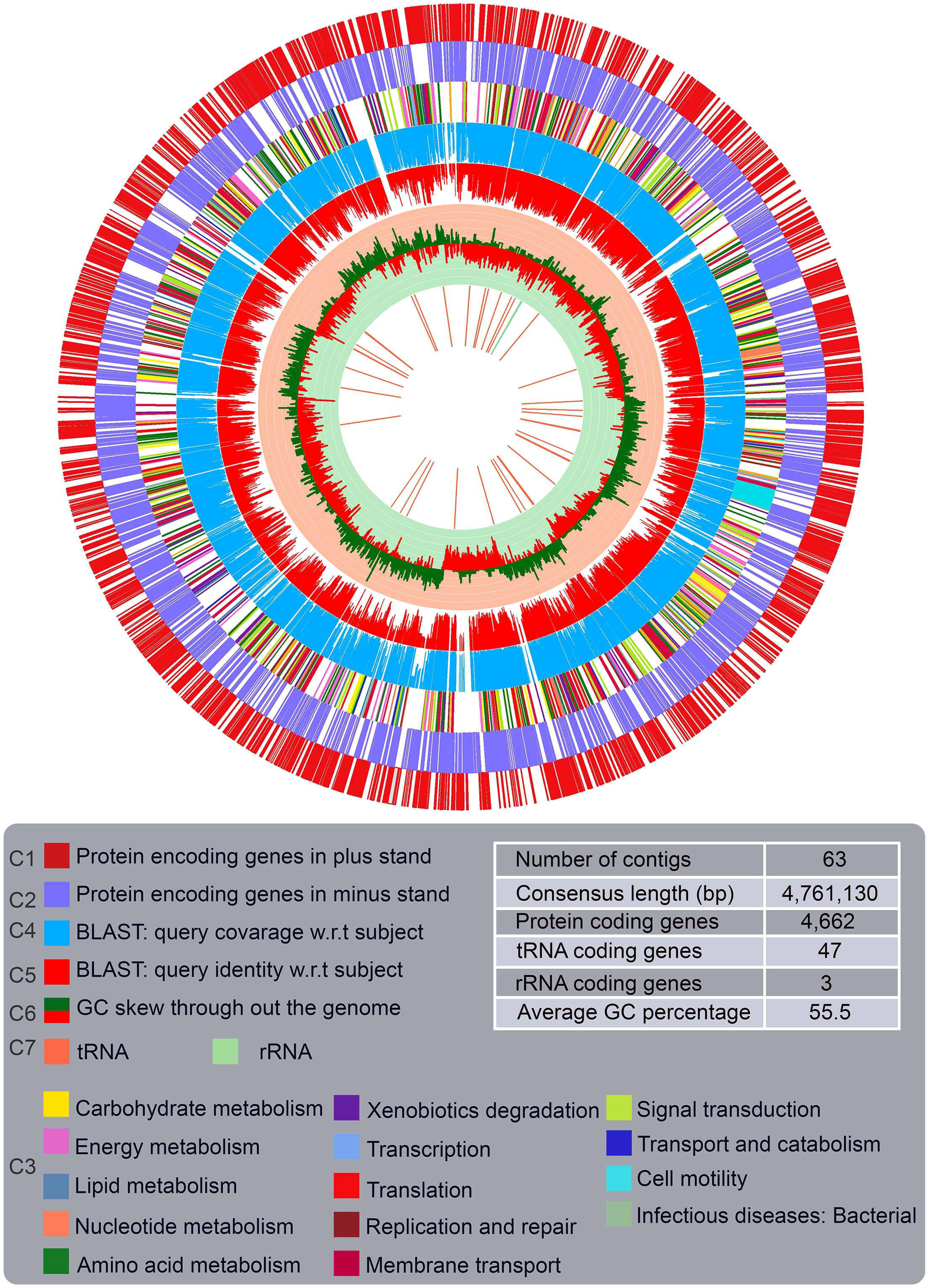
Graphical representation of the assembled and annotated whole genome shotgun sequence of *P. ginsengisoli* SBSA. The seven concentric circles, from the periphery to center represent the following: (circles 1 and 2) predicted protein-coding regions on the plus and minus strands respectively; (circle 3) protein encoding genes classified into some metabolic categories, based on KEGG orthology (KO) prediction of their products (color-codes for the metabolic categories considered are given in the lower panel); (circles 4 and 5) percentages of coverage and identity obtained respectively for all the protein encoding genes of SBSA via BlastX search of their translated amino acid sequences against the putative protein catalog derived from sequenced *Pusillimonas ginsengisoli* KCTC; (circle 6) GC skew plot embedded in light pink and sea green background; (circle 7) genes encoding homologs of known sulfur oxidation proteins, tRNAs and rRNAs. Order of arrangement, and orientation, of contigs are arbitrary, and so may not tally with the actual genome.

Apart from PGAP annotation, assembled contigs were also searched manually for homologs of genes that are reported to be coding for the relevant enzymes involved in sulfur oxidation in other bacterial members. Homologs which were detected in this way further analyzed to derive primary inferences about their impending involvement in the sulfur oxidation in *P. ginsengisoli* SBSA. A genomic locus consisting *sox* operon was detected in the genome of SBSA which encodes the components of Sox multi enzyme complex. The predicted protein products of the *sox* structural genes of the strain SBSA have highest identities with homologs from draft genome of *Pusillimonas ginsengisoli* KCTC 22046 (ID:76062; NZ_SDQE01000001.1), that have no functional evidence of sulfur-oxidation. The *sox* gene cluster in the genome of strain SBSA had the identical gene arrangement as that of the strains of *Pusillimonas ginsengisoli* KCTC 22046 and *Advenella kashmirensis* WT001 (Fig. 3.). *sox* operon of *Pusillimonas ginsengisoli* SBSA consisted genes encoding components of Sox system in the order of *soxCDYZAXOB* [(locus tags = D7I39_07160, D7I39_07165, D7I39_07170, D7I39_07175, D7I39_07180, D7I39_07185, D7I39_07190 and D7I39_07195) (Fig. 3.)]. Additionally in *P. ginsengisoli* genome, 321 amino acid (aa) long, translated product of one ORF (Protein ID-D7I39_17460) detected which shows 62% identity with *TsdA* of *A. kashmirensis* which is involved in partial conversion of thiosulfate to tetrathionate [24].

**Figure 3.**
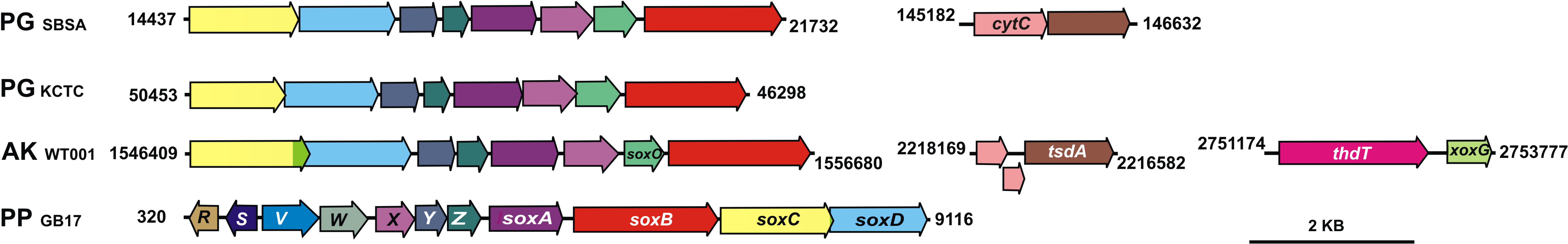
Schematic diagram illustrating syntenies of the thiosulfate/tetrathionate oxidation related genes of *P. ginsengisoli* SBSA along with its taxonomic relatives. Schematic overview of the *sox* loci of chemolithotrophic *Alphaproteobacteria P. pantotrophus* (PP) (is labeled with relevant gene names; GenBank sequence X79242.5) *P. ginsengisoli* (SBSA), *P. ginsengisoli* (KCTC) and *A. kashmirensis* (AK). Additionally *tsdA* locus of *P. ginsengisoli* SBSA and *A. kashmirensis* and *thdT* gene cluster of *A. kashmirensis* are also given. All homologs of these genes are identified by similar arrow colors. Nucleotide numbers indicate the position of the gene loci on the relevant complete genomes or partially sequenced genomic segments/contigs.

Potential promoter sequences within *P. ginsengisoli sox* and *tsdA* loci were predicted, using the web-based servers BDGP (https://www.fruitfly.org/seq_tools/promoter.html), Promoter 2.0 (http://www.cbs.dtu.dk/services/Promoter/). Two promoter-like sequences were identified within sox locus: one upstream of *soxO*, within the *soxX* ORF (indicated as P_II_ in Fig. 1A), and another in the upstream of *soxD* (indicated as P_I_ in Fig. 1A). One potential promoter sequence was predicted to be immediate upstream of *cytC* which preceded *tsdA* (indicated as P_T_ in Fig. 1A). Notably, no promoter was predicted in the region upstream of *soxC*, the first gene of this cluster.

However, no homolog of doxDA was detected in the *P. ginsengisoli* SBSA genome. Notably, no ORF similar to XoxF/ThdT [known to be involved in thiosulfate dehydrogenation and tetrathionate oxidation in *A. kashmirensis* [24] was found in the *P. ginsengisoli* SBSA genome. Within the domain Bacteria, tetrathionate hydrolase (TetH) of the *Acidithiobacillus* species [5, 17, 22] is the only tetrathionate-oxidizing enzyme that has been characterized so far at the molecular level. In this context, it is however remarkable to note that no ORF similar to *tetH* was found in the *P. ginsengisoli* genome. Other sulfur-oxidation-related genes such as reverse dissimilatory sulfite reductase and sulfide oxidase are absent in *P. ginsengisoli* SBSA; however, homologs of the sulfite: acceptor oxidoreductase genes *sorA* and *sorB* [10, 11] are present (locus tags = D7I39_01415 and D7I39_01420 respectively) in the *P. ginsengisoli* SBSA genome.

### 3.3 Involvement of *tsdA*, *soxYZ* and *soxB* in tetrathionate formation and oxidation

The *tsdA*, *soxYZ* and *soxB* genes of the *P. ginsengisoli* were individually inactivated by homologous-recombination-based insertion of kanamycin-resistance cassette (Kan^R^), resulting in the generation of the mutant strains Pg_KoT, Pg_KoYZ and Pg_KoB respectively and the sulfur-oxidizing potentials of the respective mutants were compared with that of the wild-type, by growing them in MSTy or MSTry medium (Fig. 1.). Localization/orientation of the kan^R^ cassette in the *tsdA*, *soxYZ* and *soxB* locus of Pg_KoT, Pg_KoYZ and Pg_KoB was checked by PCRs bracketing genome:*kan*^*R*^ - cassette intersection regions, followed by sequencing of the PCR products. This showed that in the mutants *kan*^*R*^ cassettes had been integrated in orientations analogous to the transcriptional direction of *tsdA*, *sox YZ* and *sox B*.

Tetrathionate oxidizing ability of Pg_KoT remained intact as observed in wild-type (however tetrathionate oxidation rate is slightly faster compared to wild type strain) when grown in MSTrY medium (Fig 1E) but in MSTY, chemolithotrophic oxidation of thiosulfate to tetrathionate was totally abolished (Fig. 1D). However, impairment of chemolithotrophic thiosulfate oxidation by Pg_koT mutant in MSTY indicates the absence of a functional Sox pathway for thiosulfate oxidation in this organism even though all the necessary genes are present in its genome. This interpretation was found to be correct when Pg_KoYZ showed physiological behavior similar to the wild type strain in MSTY and MSTrY medium respectively (Fig.1F and 1G). Tetrathionate-oxidizing abilities of the mutant Pg_KoB (Fig.1I) was totally abolished when grown in MSTrY medium. Whereas in MSTY medium, Pg_KoB converted 40mM S thiosulfate to equivalent amount of tetrathionate within 48 h of incubation (Fig.1H).

### 3.4 Substrate dependent O_2_ consumption study entails the role of glutathione in tetrathionate oxidation

Sulfur substrate dependent oxygen consumption rates of MSTY or MSTrY-growing resting cells of *P. ginsengisoli* were measured *per se*, as well as after treating them with the thiol-binding reagents N-ethyl maleimide (NEM) and iodoacetamide. Thiosulfate-dependent oxygen consumption rates of MSTY or MSTrY-grown untreated cells ranged between 90 and 110 nmol of of oxygen mg protein^−1^ min^−1^. When thiol binding reagents treated with similarly grown cells, these rates reduced to 92 nmol of oxygen mg protein^−1^ min^−1^ for NEM-treated cells and 90 nmol of oxygen mg protein^−1^ min^−1^ for iodoacetamide-treated cells (Fig. 4A). Tetrathionate-dependent oxygen consumption rates of MSTY- or MSTrY-grown untreated cells ranged between 80 and 98 nmol of oxygen mg protein^−1^ min^−1^, but cells treated with the thiol-binding reagents exhibited only 19 nmol of oxygen mg protein^−1^ min^−1^ by NEM-treated cells and 22 nmol of oxygen mg protein^−1^ min^−1^ by iodoacetamide-treated cells (Fig. 4B). Sulfite-dependent oxygen consumption rate, nonetheless, was unaffected by thiol-binding reagents (Fig. 4C). As MSTY and MSTrY-grown cells showed equivalent results for all the above experiments, data for only the former are shown (Fig. 4A–C). Reduction in tetrathionate-dependent oxygen consumption evidenced the centrality of thiol groups in tetrathionate oxidation.

**Figure 4.**
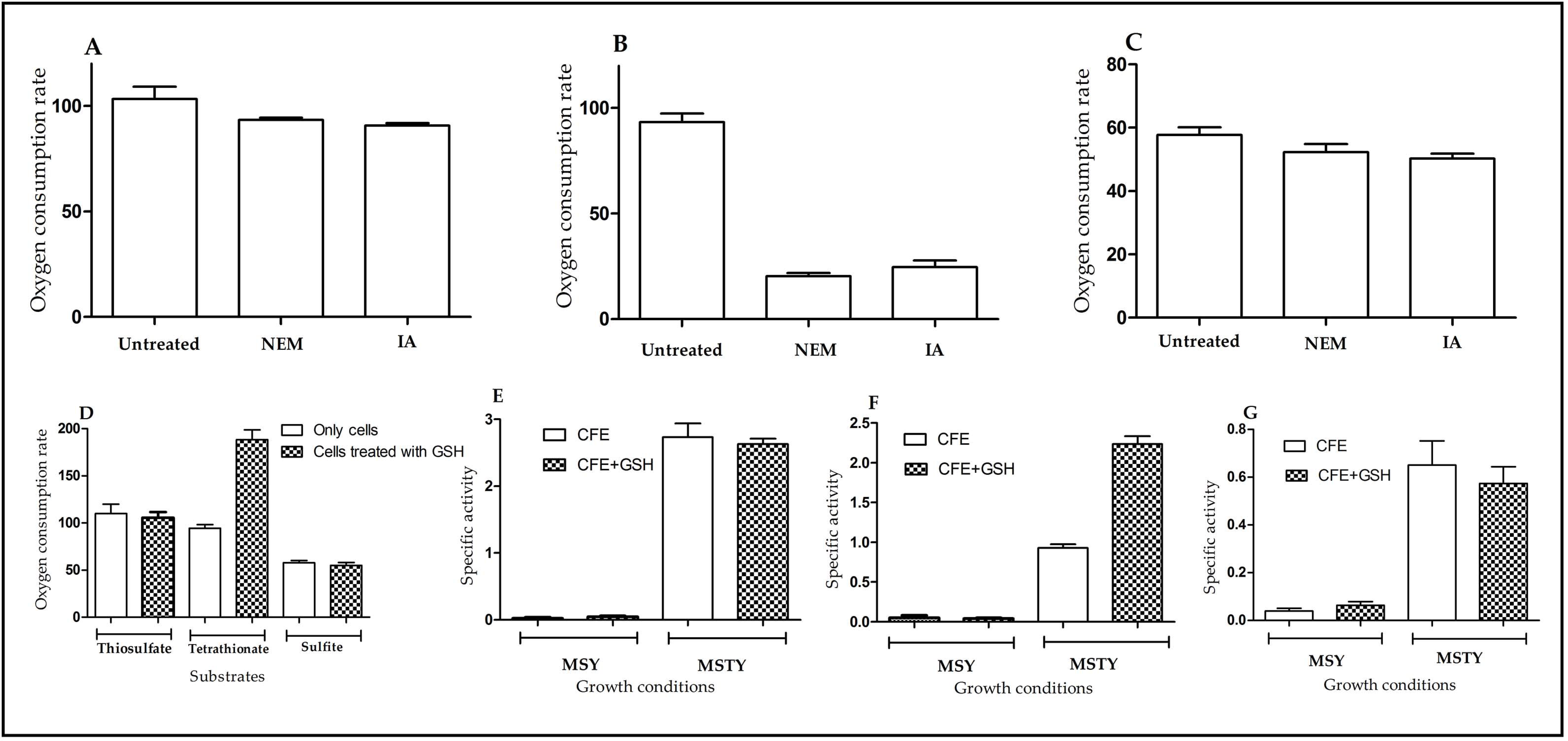
Involvement of thiols, particularly GSH, in tetrathionate oxidation by *P. ginsengisoli* SBSA. A–C. Oxygen consumption rates (expressed as nmol of oxygen consumed mg cellular protein^−1^ min^−1^) of MST-grown SBSA cells, measured in the presence of (A) thiosulfate, (B) tetrathionate or (C) sulfite, after treating with the thiol binding agent N-ethyl maleimide (NEM) or iodoacetamide (IA). Oxygen consumption rates of MSTY grown cells in the presence of the same sulfur compounds without any thiol-binding agent-treatment are given for comparison. D. Comparison of the oxygen consumption rates (in the presence of thiosulfate, tetrathionate or sulfite) of GSH-treated MSTY-grown cells with those of untreated cells. E–G. Specific activities (measured as nmol of ferricyanide reduced mg protein^−1^ min^−1^) of the (A) thiosulfate-, (B) tetrathionate or (C) sulfite-oxidizing enzymes/enzyme systems of SBSA, measured in the cell free extracts of MSY-, MSTY grown cells, in presence or absence of GSH.

Role of glutathione as the thiol-source necessary for tetrathionate oxidation was studied by comparing the substrate-dependent oxygen consumption rates of chemolithotrophically grown cells pre-incubated with GSH with that of cells not incubated with GSH. A twofold rise in the rate of tetrathionate-dependent (and not thiosulfate- or sulfite-dependent) oxygen consumption was observed for both MSTy- and MSTry-grown cells pre-incubated with 10 mM GSH in comparison to the untreated sets (Fig. 4D). Since MSTY- and MSTrY-grown cells showed equivalent rate enhancements, data for only the former types are shown (Fig. 4D). Effects of GSH on the activities of the thiosulfate-, tetrathionate and sulfite-oxidizing enzymes of *P. ginsengisoli* were also checked in the cell-free extracts of this organism. Here, also MSTY- and MSTrY-grown cells yielded equivalent results, so data for only the former types are shown in comparison with the negligible enzyme activities of MSY-grown cells (Fig. 4E–G). In the absence of GSH, specific activities of the thiosulfate, tetrathionate- and sulfite-oxidizing enzyme systems (measured as µmol ferricyanide reduced mg protein^−1^ min^−1^) were found to be 2.90, 0.98 and 0.74 respectively. The specific activity of the tetrathionate-oxidizing enzyme system increased to 2.34 when GSH was added to the reaction mixture (Fig. 4F), whereas activities of the thiosulfate- and sulfite-oxidizing systems remained more or less unchanged (Fig. 4E and G). When the substrate dependent oxygen consumption, as well as the enzyme activity, assays were repeated by substituting GSH with cysteine as the thiol source, no improvement in tetrathionate oxidation efficiency was observed.

## 4. Discussion

So far as the sulfur oxidation in *Alcaligenaceae* family is concerned, only members of *Advenella* [24], *Alcaligenes* [39], *Bordetella* [40], have the sulfur oxidation phenotype while *Achromobacter* despite having complete *sox* cluster does not have any sulfur chemotrophic attribute. However, recently one thiosulfate oxidizing species of *Pusillimonas* has been reported from activated sludge, does not harbors any homologs of structural *sox* genes in its genome [41]. To the best of our knowledge we first time bring to the fore establishment and molecular characterization of sulfur chemolithotrophy in the genus *Pusillimonas*. Present investigation revealed that the facultative sulfur chemolithotrophic strain *P. ginsengisoli* SBSA is able to utilize thiosulfate and tetrathionate as chemolithotrophic substrates. The genome of SBSA possesses a *sox* locus which shows high similarity with the *P. ginsengisoli* KCTC and *A. kashmirensis* WT001 in terms of gene synteny and their sequences. Although sox locus of SBSA contains *sox* structural genes namely *soxCDYZAXOB* but the regulatory *soxSRT* and *soxEFGH* genes are absent, products of which considered non-essential for *in vitro* thiosulfate oxidation but involved in regulation and maintenance of Sox structural proteins.

Physiological, genomic, and molecular genetic data confirmed that thiosulfate to tetrathionate conversion by *P. ginsengisoli* SBSA is governed by a homolog of TsdA. The inability of Pg_KoT to oxidize thiosulfate, also confirmed that this organism cannot render Sox mediated thiosulfate to sulfate conversion. This, however, was not unexpected considering the thiosulfate oxidation phenotype and the genetic resources for the same of its taxonomic relative [24]. Oxidation of tetrathionate to sulfate, on the other hand, involved SoxB but not SoxYZ. A similar pattern has already reported in its immediate taxonomic relative *Advenella kashmirensis* where a novel Thiol dehydrotransferase catalyzes a reaction conjugating tetrathionate to a glutathione molecule releasing one terminal sulfone sulfur from tetrathionate as sulfite which gets converted to sulfate by the action of SorAB [24]. The glutathione sulfodisulfane adduct formed in this way is oxidized to corresponding sulfates by iterative action of SoxB and SoxCD [24]. In SBSA, from the MSTY growth kinetics of Pg_KOYZ, it is confirmed that SoxYZ is not involved in tetrathionate oxidation (as in *A. kashmirensis* also). Furthermore, it is found that sox genes of SBSA always clustered as immediate sister groups of *A. kashmirensis* in phylogenetic trees constructed with individual nucleotide sequences of *soxXAYZBCD* (data not shown). Together with enhancement of tetrathionate oxidation by glutathione and presence of *sorAB* genes in SBSA, unequivocally suggest that tetrathionate oxidation in SBSA involves glutathione and downstream oxidation of tetrathionate might follow a similar mechanism as observed in *A. kashmirensis*. So the glutathione coupling with tetrathionate and downstream activity of SoxB, SoxCD, and SorAB appear to be instrumental in tetrathionate oxidation by SBSA but we could not detect a ThdT like protein as there was no genome-based clue to suspect any protein known to be involved in this process. That will be a next generation of research to find out the potential proteins involved in glutathione-tetrathionate coupling and its downstream oxidation in this organism/ organism with similar sulfur oxidation phenotype and genetic repository. Furthermore, as the Pg_KoT unable to convert thiosulfate to tetrathionate, it can be assumed that novel protein(s) involved in glutathione-tetrathionate coupling in SBSA, is not a dual functional protein like ThdT.

During the chemolithotrophic oxidation of thiosulfate by SBSA, tetrathionate and sulfate formation start simultaneously, thereby forms equivalent fractions of tetrathionate and sulfate until the thiosulfate is exhausted completely in the spent medium. This may be due to similar substrate-dependent oxygen consumption rate of thiosulfate and tetrathionate.

The genetic basis of thiosulfate oxidation by facultative chemolithotrophic members of the class beta proteobacteria is experimentally studied only in *P. ginsengisoli* SBSA and *A. kashmirensis* WT001 till date. These two organisms oxidize intermediary formed tetrathionate with the involvement of SoxB, SoxCD but not SoxYZ. Both these organisms despite having *sox* clusters in their genomes, do not oxidize thiosulfate directly to sulfate. This non-fucntional status of their *sox* genes could have stemmed from their alternative strategy for thiosulfate oxidation via S_4_I pathway. Furthermore, it is also noteworthy that a single gene-single phenotype of TsdA have obvious advantage over multiple gene-coded Sox mechanism in respect to both horizontal gene transfer mediated gain / loss of function via random mutagenesis. Based on these observations it could be hypothesized that in due course of evolution, *sox* genes that are involved in tetrathionate oxidation remain functional while those do not (like *soxYZ* and *soxXA*) may become vestigial components of the genome particularly where catalytically strong TsdA is present as an alternative S_4_I strategy for thiosulfate oxidation.

## Acknowledgements

This research was funded by Bose Institute, Department of Science and Technology, Government of India (GOI). SM got fellowship from Department of Science and Technology, GoI. MJR got fellowship from University Grants Commission. JS and PP received fellowship from the Council of Scientific and Industrial Research (CSIR), S.B. received fellowship from Bose Institute.

## Supplementary data

Supplemental material for this article may be found with the digital version of this manuscript.

## References

[1] Ghosh W, Dam B. Biochemistry and molecular biology of lithotrophic sulfur oxidation by taxonomically and ecologically diverse bacteria and archaea. FEMS Microbiol. Rev. 2009; 33:999–1043.

[2] Liu Y, Beer LL, Whitman WB. Sulfur metabolism in archaea reveals novel processes. Environ. Microbiol. 2012; 14:2632–2644.

[3] Friedrich CG, Rother D, Bardischewsky F, Quentmeier A, Fischer J. Oxidation of reduced inorganic sulfur compounds by bacteria: emergence of a common mechanism?. Appl. Environ. Microbiol. 2001; 67:2873–82.

[4] Müller FH, Bandeiras TM, Urich T, Teixeira M, Gomes CM, Kletzin A. Coupling of the pathway of sulphur oxidation to dioxygen reduction: characterization of a novel membrane-bound thiosulphate: quinone oxidoreductase. Molecular microbiology. 2004; 53:1147–60.

[5] Rzhepishevska OI, Valdés J, Marcinkeviciene L, Gallardo CA, Meskys R, Bonnefoy V, Holmes DS, Dopson M. Regulation of a novel *Acidithiobacillus caldus* gene cluster involved in metabolism of reduced inorganic sulfur compounds. Appl. Environ. Microbiol. 2007; 73:7367–72.

[6] Kletzin A. Oxidation of sulfur and inorganic sulfur compounds in *Acidianus ambivalens*. In Microbial sulfur metabolism 2008; 184–201. Springer, Berlin, Heidelberg.

[7] Liu LJ, Stockdreher Y, Koch T, Sun ST, Fan Z, Josten M, et al. Thiosulfate transfer mediated by DsrE/TusA homologs from acidothermophilic sulfur-oxidizing archaeon Metallosphaera cuprina. Journal of Biological Chemistry. 2014; 289:26949–59.

[8] Kelly DP, Shergill JK, Lu WP, Wood AP. Oxidative metabolism of inorganic sulfur compounds by bacteria. Antonie Van Leeuwenhoek. 1997; 71:95–107.

[9] Appia-Ayme C, Little PJ, Matsumoto Y, Leech AP, Berks BC. Cytochrome complex essential for photosynthetic oxidation of both thiosulfate and sulfide in Rhodovulum sulfidophilum. J. bacterial. 2001; 183:6107–18.

[10] Kappler U, Friedrich CG, Trüper HG, Dahl C. Evidence for two pathways of thiosulfate oxidation in *Starkeya novella* (formerly *Thiobacillus novellus*). Archives of Microbiology. 2001;175:102–11.

[11] Kappler U, Aguey-Zinsou KF, Hanson GR., Bernhardt PV, McEwan AG. Cytochrome c551 from *Starkeya novella* characterization, spectroscopic properties, and phylogeny of a diheme protein of the SoxAX family. J Biol Chem. 2004; 279: 6252–6260.

[12] Mukhopadhyaya PN, Deb C, Lahiri C, Roy P. A soxA gene, encoding a diheme cytochrome c, and a sox locus, essential for sulfur oxidation in a new sulfur lithotrophic bacterium. J. bacterial. 2000; 182:4278–87.

[13] Bamford VA, Bruno S, Rasmussen T, Appia-Ayme C, Cheesman MR, Berks BC, Hemmings AM. Structural basis for the oxidation of thiosulfate by a sulfur cycle enzyme. The EMBO journal. 2002; 21:5599–610.

[14] Sauvé V, Bruno S, Berks BC, Hemmings AM. The SoxYZ complex carries sulfur cycle intermediates on a peptide swinging arm. J Biol Chem. 2007; 282:23194–204.

[15] Boden R. Reclassification of Halothiobacillus hydrothermalis and Halothiobacillus halophilus to Guyparkeria gen. nov. in the Thioalkalibacteraceae fam. nov., with emended descriptions of the genus Halothiobacillus and family Halothiobacillaceae. Int J Syst Evol Microbiol. 2017; 67:3919–3928.

[16] Visser JM, de Jong GA, Robertson LA, Kuenen JG. Purification and characterization of a periplasmic thiosulfate dehydrogenase from the obligately autotrophicThiobacillus sp. W5. Arch Microbiol. 1996; 166:372–8.

[17] De Jong GA, Hazeu W, Bos P, Kuenen JG. Isolation of the tetrathionate hydrolase from Thiobacillus acidophilus. Eur J Biochem. 1997; 243:678–83.

[18] Bugaytsova Z, Lindström EB. Localization, purification and properties of a tetrathionate hydrolase from Acidithiobacillus caldus. Eur J Biochem. 2004; 271:272–80.

[19] Ghosh W, Bagchi A, Mandal S, Dam B, Roy P. Tetrathiobacter kashmirensis gen. nov., sp. nov., a novel mesophilic, neutrophilic, tetrathionate-oxidizing, facultatively chemolithotrophic betaproteobacterium isolated from soil from a temperate orchard in Jammu and Kashmir, India. India. Int J Syst Evol Microbiol. 2005; 55:1779–87.

[20] Dam B, Mandal S, Ghosh W, Gupta SK, Roy P. The S4-intermediate pathway for the oxidation of thiosulfate by the chemolithoautotroph *Tetrathiobacter kashmirensis* and inhibition of tetrathionate oxidation by sulfite. Res Microbiol. 2007; 158:330–8.

[21] Kikumoto M, Nogami S, Kanao T, Takada J, Kamimura K. Tetrathionate-forming thiosulfate dehydrogenase from the acidophilic, chemolithoautotrophic bacterium *Acidithiobacillus ferrooxidans*. Appl. Environ. Microbiol. 2013; 79:113–20.

[22] Kanao T, Kamimura K, Sugio T. Identification of a gene encoding a tetrathionate hydrolase in *Acidithiobacillus ferrooxidans*. J. Biotechnol. 2007; 132:16–22.

[23] Quatrini R, Appia-Ayme C, Denis Y, Jedlicki E, Holmes DS, Bonnefoy V. Extending the models for iron and sulfur oxidation in the extreme acidophile. Acidithiobacillus ferrooxidans. BMC genomics. 2009; 10:394.

[24] Pyne P, Alam M, Rameez MJ, Mandal S, Sar A, Mondal N et al. Homologs from sulfur oxidation (Sox) and methanol dehydrogenation (Xox) enzyme systems collaborate to give rise to a novel pathway of chemolithotrophic tetrathionate oxidation. Mol. Microbiol. 2018; 109:169–91.

[25] Hensen D, Sperling D, Trüper HG, Brune DC, Dahl C. Thiosulphate oxidation in the phototrophic sulphur bacterium Allochromatium vinosum.. Mol. Microbiol. 2006; 62:794–810.

[26] Brito JA, Denkmann K, Pereira IA, Archer M, Dahl C. Thiosulfate Dehydrogenase (TsdA) from *Allochromatium vinosum* structural and functional insights into thiosulfate oxidation. J Biol Chem. 2015; 290:9222–38.

[27] Fernandes S, Mazumdar A, Bhattacharya S, Peketi A, Mapder T, Roy R et al. Enhanced carbon-sulfur cycling in the sediments of Arabian Sea oxygen minimum zone center. Sci Rep. 2018; 8:8665.

[28] Ghosh W, Roy P. Chemolithoautotrophic oxidation of thiosulfate, tetrathionate and thiocyanate by a novel rhizobacterium belonging to the genus Paracoccus. FEMS Microbiol Lett. 2007; 270:124–31.

[29] Euzéby JP. List of Bacterial Names with Standing in Nomenclature: a folder available on the Internet. Int J Syst Evol Microbiol. 1997; 47:590–2.

[30] Parte AC. LPSN—list of prokaryotic names with standing in nomenclature. Nucleic Acids Res. 2013; 42:613–D616.

[31] Kelly DP, Wood AP. Synthesis and determination of thiosulfate and polythionates. In Methods Enzymol. 1994 Jan 1 (Vol. 243, pp. 475–501). Academic Press.

[32] Alam M, Pyne P, Mazumdar A, Peketi A, Ghosh W. Kinetic enrichment of 34S during proteobacterial thiosulfate oxidation and the conserved role of SoxB in SS bond breaking. Appl. Environ. Microbiol.. 2013; 79:4455–4464.

[33] West PW, Gaeke GC. Fixation of sulfur dioxide as disulfitomercurate (II) and subsequent colorimetric estimation. Anal Chem. 1956; 28: 1816–1819.

[34] Hilger D, Jung H. Protein Chemical and Electron Paramagnetic Resonance Spectroscopic Approaches to Monitor Membrane Protein Structure and Dynamics–Methods Chapter. Bacterial Signaling. 2009: 247–263.

[35] Trudinger PA. Thiosulphate oxidation and cytochromes in Thiobacillus X. 1. Fractionation of bacterial extracts and properties of cytochromes. Biochem J. 1961; 78:673.

[36] Bankevich A, Nurk S, Antipov D, Gurevich AA, Dvorkin M, Kulikov AS et al. SPAdes: a new genome assembly algorithm and its applications to single-cell sequencing. J Comput Biol. 2012; 19:455–477.

[37] Parks DH, Imelfort M, Skennerton CT, Hugenholtz P, Tyson GW. CheckM: assessing the quality of microbial genomes recovered from isolates, single cells, and metagenomes. Genome Res. 2015; 25:1043–1055.

[38] Kurtz S, Phillippy A, Delcher AL, Smoot M, Shumway M, Antonescu C et al. Versatile and open software for comparing large genomes. Genome Biol. 2004; 5:12.

[39] Aguilar JR, Cabriales JJ, Vega MM. Identification and characterization of sulfur-oxidizing bacteria in an artificial wetland that treats wastewater from a tannery. Int J Phytoremediation. 2008; 10:359–70.

[40] Nisola GM, Tuuguu E, Farnazo DM, Han M, Kim Y, Cho E et al. Hydrogen sulfide degradation characteristics of Bordetella sp. Sulf-8 in a biotrickling filter. Bioprocess and biosystems engineering. 2010; 33:1131–8.

[41] Koh HW, Song MS, Do KT, Kim H, Park SJ. *Pusillimonas thiosulfatoxidans* sp. nov., a thiosulfate oxidizer isolated from activated sludge. Int J Syst Evol Microbiol. 2019; 69:1041–6.

[42] Simon RU, Priefer U, Pühler A. A broad host range mobilization system for in vivo genetic engineering: transposon mutagenesis in gram negative bacteria. Biotechnology. 1983; 1:784.

[43] Skorupski K, Taylor RK. Positive selection vectors for allelic exchange. Gene. 1996; 169:47–52.

