## Supplementary material for "Molecular mechanism of sulfur chemolithotrophy in the betaproteobacterium *Pusillimonas ginsengisoli*"

Running Title: Sulfur-chemolithotrophic pathway of *Pusillimonas ginsengisoli*

**Contents**

**Supporting experimental procedures:**

I. Knock-out mutagenesis: mutation of *tsdA* gene.
II. Knock-out mutagenesis: mutation of *soxYZ*.
III. Knock-out mutagenesis: mutation of *soxB*.

**Supplementary tables**

1. Tables S1 through S3, with relevant titles and notes.

***Knock-out mutagenesis: mutation of tsdA***

The *tsdA* gene of the *P. ginsengisoli* sox operon did not contain any unique restriction site where an antibiotic-resistance cartridge could be inserted. So a unique Kpn I site was incorporated between nucleotide positions 1517 and 1523 of this gene via fusion PCR method (Shevchuk et al., 2004). Two PCR reactions were carried out using the primer pairs Pg_KOtsd_F and Pg_KOtsd_FU_R, and Pg_KOtsd_FU_F and Pg_KOtsd_R respectively (see supplementary Table S1). This resulted in two 1047 and 1022 bp PCR products respectively. The primers Pg_KOtsd_FU_R and Pg_KOtsd_FU_F were designed in such a way that each binds in the same position of the gene but with the opposite strands and in reverse orientation, even as each possess an KpnI restriction site in the same position. The final round of PCR, where a mixture of purified FUT1 and FUT2 was used as template and Pg_KOtsd_F and Pg_KOtsd_R were used as primers, produced a 2096 bp amplicon containing an KpnI site inside the *tsdA* allele. This PCR product was cloned into pKAS32 between the XbaI and SacI sites, which gave rise to the plasmid pGKotsd. On the other hand, the Kan-resistance (*KanR*) gene with its associated promoter was amplified from pUC4K plasmid by using the primer pair kanR_KpnI_F and kanR_KpnI_R (see supplementary Table S1). The kanR PCR product (1136 bp) was then inserted within the KpnI site of the recombinant pGKotsd which gave rise to the final construct pGKotsd::KanR, which in its turn was transformed into *E. coli* S17.1 to produce the donor strain and finally delivered into *P. ginsengisoli* SBSA_SR from the donor *E. coli* S17.1 via chemical transformation method.

After the purported first cross-over, colonies having the phenotype KanR (Kanmycin-resistant), AmpR (Ampicilin resistant) and SmS (Streptomycin-sensitive) were selected as single crossover mutants. These were then sub-cultured on LA plates supplemented with Sm (100 µg ml^-1^) and Kan (40 µg ml^-1^) to obtain double crossover mutants that had apparently lost the plasmid backbone (pGKotsd::rpsl) and retained only the tsdA -integrated kanR gene. Double crossover mutants were further checked by PCR with the primer pair Pg_KOtsd_Internal_F and Pg_KOtsd_Internal_R and sequencing of the obtained products.

***Knock-out mutagenesis: mutation of soxYZ***

Like *tsdA* gene, *soxYZ* gene of the *P. ginsengisoli* sox operon did not contain any unique restriction site, so a unique Kpn I site was incorporated between nucleotide positions 851 and 857 of this gene via fusion PCR method. Two PCR reactions were carried out using the primer pairs Pg_KOYZ_F and Pg_KOYZ_FU_R, and Pg_KOYZ_FU_F and Pg_KOYZ_R respectively (see Supporting Table S2). This resulted in two 1069 and 1064 bp PCR products, viz. FUY1 and FUY2 respectively. The primers Pg_KOYZ_FU_R and Pg_KOYZ_FU_F were designed in such a way that each binds in the same position of the gene but with the opposite strands and in reverse orientation, even as each possess an KpnI restriction site in the same position. The final round of PCR, where a mixture of purified FUY1 and FUY2 was used as template and Pg_KOYZ_F and Pg_KOYZ_R were used as primers, produced a 2137 bp amplicon containing an KpnI site inside the *soxYZ* allele. This PCR product was cloned into pKAS32 between the XbaI and SacI sites, which gave rise to the plasmid pGKoYZ. On the other hand, the Kan-resistance (*KanR*) gene with its associated promoter was amplified from pUC4K plasmid by using the primer pair kanR_KpnI_F and kanR_KpnI_R (see Supporting Table S2). The kanR PCR product was then inserted within the KpnI site of the recombinant pGKoYZ which gave rise to the final construct pGKoYZ::KanR, which in turn was transformed into *E. coli* S17.1 to produce the donor strain and finally delivered into *P. ginsengisoli* SBSA_SR from the donor *E. coli* S17.1 via chemical transformation method. Double crossover mutants were further checked by PCR with the primer pair Pg_KOYZ_Internal_F and Pg_KOYZ_Internal_R and sequencing of the obtained products following similar method used for *tsdA* gene.

***Knock-out mutagenesis: mutation of soxB***

Unique Kpn I site was incorporated between nucleotide positions 1100 and 1106 of the *soxB* gene via fusion PCR method as it did not contain any unique restriction site. Two PCR reactions were carried out using the primer pairs Pg_KOB_F and Pg_KOB_FU_R, and Pg_KOB_FU_F and Pg_KOB_R respectively (see Supporting Table S3). This resulted in two 1054 and 1084 bp PCR products, viz. FUB1 and FUB2 respectively. The primers Pg_KOB_FU_R and Pg_KOB_FU_F were designed in such a way that each binds in the same position of the gene but with the opposite strands and in reverse orientation, even as each possess an KpnI restriction site in the same position. The final round of PCR, where a mixture of purified FUB1 and FUB2 was used as template and Pg_KOB_F and Pg_KOB_R were used as primers, produced a 2138 bp amplicon containing an KpnI site inside the *soxB* allele. This PCR product was cloned into pKAS32 between the XbaI and SacI sites, which gave rise to the plasmid pGKoB. On the other hand, the Kan-resistance (*KanR*) gene with its associated promoter was amplified from pUC4K plasmid by using the primer pair kanR_KpnI_F and kanR_KpnI_R (see Supporting Table S3). The kanR PCR product was then inserted within the KpnI site of the recombinant pGKoB which gave rise to the final construct pGKoB::KanR, which in its turn was transformed into *E. coli* S17.1 to produce the donor strain and finally delivered into *P. ginsengisoli* SBSA_SR from the donor *E. coli* S17.1 via chemical transformation method. Double crossover mutants were further checked by PCR with the primer pair Pg_KOB_Internal_F and Pg_KOB_Internal_R and sequencing of the obtained products.

**Table S1: Primers used in the knock-out mutagenesis of *tsdA* gene of *P. ginsengisoli*. The XbaI site is in bold while SacI is bold and underlined font, the Kpn I sites is underlined.**

| **Sl.No.** | **Primer Name** | **Primer sequence (5’-3’)** |
| --- | --- | --- |
| 1 | Pg_KOtsd_F | AGCG**TCTAGA**ACGTTGCCCCTATTGACACC |
| 2 | Pg_Kotsd_FU_R | TTCACAGACTCGGGCAGCCCTGAAAAATAG |
| 3 | Pg_KO_tsd_FU_F | CTATTTTTCAGGGCTGCCCGAGTCTGTGAACGAGGGTACCCATGATTGATGCACTGATAT |
| 4 | Pg_Kotsd_R | ACTA**GAGCTC**GTAAAGCAGGCCCTGGACTT |
| 5 | Pg_KOtsdA_Internal_F | CGCGCACTCACCCACTTACCCCGAC |
| 6 | Pg_KOtsdA_Internal_R | TGATGAATGCCGCCGCTGTCGACAC |
| 7 | Pg_KOtsdA_Internal seq_F | CGACTGGGATAACAATATACCGGCC |
| 8 | Pg_KOtsdA_Internal seq_R | GCAACCTCCATGGCTTTACGCTTG |
| 9 | kanR_KpnI_F | TAGACGGTACCGTTTTATGGACAGCAAGCG |
| 10 | kanR_KpnI_R | GGGGTACCAAGAACTCCAGCATGAGATCCC |

**Table S2: Primers used in the knock-out mutagenesis of *soxYZ* gene of *P. ginsengisoli* SBSA. The XbaI site is in bold while SacI is bold and underlined font, the Kpn I sites is underlined.**

| **Sl.No.** | **Primer Name** | **Primer sequence (5’-3’)** |
| --- | --- | --- |
| 1 | Pg_KOYZ_F | GGCA**TCTAGA**CGAAAGCGGAGAGGTTTTCA |
| 2 | Pg_KOYZ_FU_R | TGCATCATCCTGGTCTTTGTACTTGGCAGC |
| 3 | Pg_KOYZ_FU_F | GCTGCCAAGTACAAAGACCAGGATGATGCAAGAAGGTACCAGTCATACGCATCCACCAGG |
| 4 | Pg_KOYZ_R | TGCA**GAGCTC**ACTGATCCAGCCTGTCCATG |
| 5 | Pg_KOYZ_Internal_F | AGCCAAAATTCAGAAGGGAGGATCCGGC |
| 6 | Pg_KOYZ_Internal_R | CCCCATTGGGCAGACAGCACAGTCTTATC |
| 7 | Pg_KOYZ_Internal seq_F | ACACGTCTGGGAAACCTGCTTCGATTCTTC |
| 8 | Pg_KOYZ_Internal seq_R | TTGGGTAGATCCTGCAACCGTATCCGCTG |
| 9 | kanR_KpnI_F | TAGACGGTACCGTTTTATGGACAGCAAGCG |
| 10 | kanR_KpnI_R | GGGGTACCAAGAACTCCAGCATGAGATCCC |

**Table S3: Primers used in the knock-out mutagenesis of *soxB* gene of *P. ginsengisoli***. The XbaI site is in bold while SacI is bold and underlined font, the Kpn I sites is underlined.

| **Sl.No.** | **Primer Name** | **Primer sequence (5’-3’)** |
| --- | --- | --- |
| 1 | Pg_KOB_F | GCTG**TCTAGA**ATGTGGCGCTGCCTGAAGTT |
| 2 | Pg_KOB_FU_R | TTCGACGCGTTCCTGTCCCAGTGTGAACTC |
| 3 | Pg_KOB_FU_F | GAGTTCACACTGGGACAGGAACGCGTCGAACGTGGGTACCGAGACAGAAAAACCATAGGC |
| 4 | Pg_KOB_R | TACG**GAGCTC**GCTGGTTCATAGAGTGCC |
| 5 | Pg_KOB_Internal_F | ATGCGCCGGGGGTTGAACTCATTGC |
| 6 | Pg_KOB_Internal_R | CAGCGTCTTGCCTCCGGCATTCTCC |
| 7 | Pg_KOB_Internal seq_F | GCTAAACCGGCCACCCCATCTG |
| 8 | Pg_KOB_Internal seq_R | CTGCGTGCGCAACAGCTACGATC |
| 9 | kanR_KpnI_F | TAGACGGTACCGTTTTATGGACAGCAAGCG |
| 10 | kanR_KpnI_R | GGGGTACCAAGAACTCCAGCATGAGATCCC |

**Reference**

Shevchuk, N.A., Bryksin, A.V., Nusinovich, Y.A., Cabello, F.C., Sutherland, M., and Ladisch, S. (2004) Construction of long DNA molecules using long PCR-based fusion of several fragments simultaneously. Nucleic Acids Res 32: 19e.

Partial sequence alignment of TsdA homologues protein from proteobacteria. Haem binding motif are indicated by red colour. Putative distal haem ligands are marked in green colour. Strictly conserved residues marked in pink colour. Species *Allocromatium vinosum* (AV), *Advenella kashmirensis* AK. *Pusillimonas ginsengisoli*, PG; *Thermithiobacillus tepederius*(SMMA18)

**Role of conserved Cys123:**The two cysteines in typical Cys-X2-Cys-His haem binding motifs of c-type cytochromes like Alc. vinosum TsdA provide the thioether linkage to the haem, and the adjacent histidine serves as the fifth ligand to the haem iron. On the opposite side of this proximal ligand a sixth ligand, the distal axial ligand, can additionally ligate the haem iron. leading to an octahedral complex. The sixth ligand is usually a histidine or methionine. While comparative sequence analysis does not indicate any conserved histidines in TsdA apart from those in the haem binding motifs, two conserved methionines (Met222 or Met236, according to Alc. vinosum numbering) are present that could act as distal axial haem ligands (Fig. 1). Most noteworthy, a strictly conserved cysteine (Cys123) is also present. It should be pointed out that His/Cys axial ligation of haem in c-type cytochromes has been described before albeit in only a few cases so far (Cheesman et al., 2001;Grein et al., 2010).
